# Decomposition in mixed beech forests in the south-western Alps under severe summer drought

**DOI:** 10.1101/2020.05.23.111815

**Authors:** Marion Jourdan, Stephan Hättenschwiler

**Affiliations:** CEFE, Univ. Montpellier, CNRS, EPHE, IRD, Univ Paul Valéry Montpellier 3, Montpellier, France; Agence de l’environnement et de la Maîtrise de l’Energie, Angers, France; ESE, CNRS, AgroParisTech, Univ. Paris-Saclay, F-91400 Orsay, France

**Keywords:** carbon cycling, French Alps, nitrogen cycling, North-South gradient, plant diversity, rainfall exclusion, reduced precipitation

## Abstract

Climate and plant litter diversity are major determinants of carbon (C) and nitrogen (N) cycling rates during decomposition. Yet, how these processes will be modified with combined changes in climate and biodiversity is poorly understood. With a multisite field experiment, we studied the interactive effects of summer drought (using rainout shelters) and tree species mixing in beech forests in the French Alps. Forests included monospecific stands of *Fagus sylvatica, Abies alba*, and *Quercus pubescens* and two-species mixtures composed of beech and one of the other species. We hypothesized (1) negative effects of summer drought on C and N loss during decomposition and (2) mitigation of these negative effects in mixed tree species stands. Litter lost 35% of initial C, and 15% of N on average across all sites and litter types over 30 months of decomposition. Summer drought consistently, but weakly, reduced C loss but had no effect on N loss. Tree species mixing did not alter drought effects on decomposition but had non-additive effects on C and N loss, which were dominated by direct litter mixing rather than indirect tree canopy effects. Our data suggest relatively small drought effects on decomposition, possibly because process rates are generally slow during summer and because microsite variability exceeds that in response to rain exclusion. The dominant contribution of litter mixing to biodiversity effects supports the importance of microsite conditions for C and N dynamics during decomposition, which should be accounted for more explicitly in climate and biodiversity change predictions.

## 1 INTRODUCTION

The general global trend of increasing temperatures since the 1980s is accompanied by higher frequencies of extreme climatic events with the potential to profoundly change ecosystem functioning (IPCC, 2014). For example, drought events become more frequent and severe (Dale and others 2001; Pachauri and others 2015) and are predicted to further increase, both in frequency and intensity during the course of the 21^st^ century (IPCC Pachauri and others 2015). These predictions vary strongly among different geographical regions (Polade and others 2017). In the Mediterranean area and the western parts of the Alps, for example, regional climate models predict a decrease in the total amount of precipitation and an increase in the duration and frequency of summer drought (Giorgi and Lionello 2008; Dubrovský and others 2014; Polade and others 2017). Because the amount and distribution of precipitation is a key climatic driver of species distribution patterns and controls ecosystem processes largely, these changes are expected to modify the composition of plant, animal and microbial communities and the functioning of ecosystems.

Decomposition contributes to the regulation of carbon (C) and nutrient cycling and is a major ecosystem process that is sensitive to climate change. Indeed, the climate variables temperature and humidity have a major control on the rates of decomposition (Berg and others 1993; Couteaux and others 1995; Berg 2000; Bradford and others 2016). Temperature and humidity determine the activity of decomposer organisms and the biochemical processes involved in the breakdown and transformation of organic matter and the mineralization of C and nutrients (Pastor and Post 1986; Aerts 1997; Sanaullah and others 2012). There are essentially two ways to experimentally assess the response of decomposition to changing climatic conditions. The space for time substitution along climatic gradients allows to quantify decomposition dynamics in an undisturbed natural context (Harmon and others 2009). Such studies laid the ground for the current understanding of climate control over decomposition (Berg and others 1993; Parton and others 2007; Bradford and others 2016). Because multiple other factors, including soil parameters, plant community composition, and the abundance and diversity of decomposer communities change alongside with climate across these gradients, it is often difficult to determine the driving mechanisms and to distinguish clearly among the different factors. A second approach is to experimentally modify the climatic conditions, for example by altering the amount of precipitation using rainfall exclusion (Yahdjian and Sala 2006; Sanaullah and others 2012; Santonja and others 2017, 2019; Pereira and others 2019). Experimental rain exclusion is logistically challenging and thus mostly restricted to individual sites within a specific bioclimatic context. The technical solutions vary considerably among studies, which makes it difficult to compare across larger spatial scales. Some field experiments use movable roofs that allow simulating predictions with variable seasonal changes, including complete rainfall exclusion for a part of the year (e.g. Santonja and others 2017). However, it is not always possible to install technically complex structures, especially in tall vegetation such as forests. Most of the experiments manipulating precipitation adopt continuous partial rain exclusion resulting in a variable percentage of reduced total rainfall over the whole year (e.g. Yahdjian and Sala 2006; García-Palacios and others 2013; Pereira and others 2019). A continuous fixed percentage of rainfall removal, however, does not necessarily match with the predictions of climate change models expecting rather different distribution patterns with extreme events, such as more intense summer drought (Giorgi and Lionello 2008; Dubrovský and others 2014; Polade and others 2017). Previous studies reported generally slower decomposition when precipitation was reduced with variable effect sizes depending on the study sites. The methods used, with typically stronger effects when rainfall was completely excluded during part of the year (Saura-Mas and others 2012; Santonja and others 2017) than with continuous partial rainfall exclusion (García-Palacios and others 2013; Almagro and others 2015; Santonja and others 2019). The consequences of extended drought periods by totally excluding rainfall on decomposer organisms and the processes they drive may differ and be stronger compared to partial but continuous reduction of rainfall (Fierer and others 2003; Schimel and others 2007). Accordingly, it was shown that microbial driven decomposition (Joly and others 2017a) and isopod effects on decomposition (Joly and others 2019) depended more strongly on the frequency than the total cumulative amount of rainfall in a Chihauhuan Desert shrubland.

It has recently been stressed that decomposition is very sensitive to variation in microclimatic conditions at very small spatial scales that can even override macroclimatic control over decomposition (Bradford and others 2016). This reconsideration of the role of macroclimatic parameters put more emphasis on local scale conditions and plant species-specific litter quality that can account for more of the variance in decomposition rates than continental scale differences in macroclimatic conditions (Cornwell and others 2008). Across a continental gradient in Europe Joly and others (2017b) showed that plant canopy characteristics predict variation in decomposition better than the large differences in macroclimate. Plant canopy composition and diversity also affected decomposition indirectly via modified environmental conditions in a plant diversity experiment (Hector 1999). Beyond these indirect plant canopy effects on the decomposer environment, the structure and diversity of the litter layer can also affect microclimatic conditions and decomposer activity (Hättenschwiler and others 2005). The different sizes, shapes and morphologies of leaf litter originating from different plant species affect the geometric organization of the litter layer and its water retention capacity, which ultimately determines microclimate and habitat structure for decomposer communities. For example, Wardle and others (2003) observed that the presence of feather mosses in litter mixtures of subarctic plant species systematically increased litter humidity through their high water retention capacity leading to higher decomposition of mixtures including feather mosses compared to mixtures without feather mosses. Similarly, Makkonen and others (2013) showed that increasing differences in water-holding capacity among distinct litter types led to more synergistic effects in litter mixture decomposition under limiting water conditions. The consequences of changing patterns in precipitation on decomposition may therefore depend to a certain degree on plant canopy characteristics, litter traits, and other local conditions. However, it is currently not well understood how the consequences of reduced precipitation on decomposition may depend on plant community characteristics. Only very few studies addressed this question at a small local scale of a particular study site (Vogel and others 2013; Santonja and others 2017) and reported that the effects of plant community composition and diversity are as important, or even more important than differences in precipitation in controlling litter decomposition. Santonja and others (2017) also found that higher litter diversity attenuated the overall negative drought effects on decomposition. How such interactions between plant community composition and changing precipitation play out at larger regional scales including different site conditions and plant communities is unknown.

Here we aimed at testing whether carbon (C) and nitrogen (N) fluxes during leaf litter decomposition are affected by extended summer drought, and whether these effects are modified by canopy composition along a north-south gradient of mixed European beech (*Fagus sylvatica*) forests in the south-west part of the French Alps. We have chosen these forests because European beech is a dominant species in temperate forests of Western Europe, and because beech seems to be particularly sensitive to a drier climate (Niinemets and Valladares 2006). Moreover, the north-south gradient implies a natural gradient in climatic conditions with generally longer and more pronounced summer drought periods in the south, which is also accompanied by a change in the co-dominant tree species associated with European beech, with silver fir (*Abies alba*) in the northern part and downy oak (*Quercus pubescens*) in the southern part. By including pure stands of each of the species and their two-species mixtures (beech-fir in the North and beech-oak in the South), these naturally occurring forests provided an ideal setup to test our hypotheses. We predicted that (1) increased summer drought has negative effects on C and N loss during decomposition, and that these negative effects are stronger in the northern sites with relatively wet summers compared to the southern sites with already quite dry summers. We also expected that (2) mixed species canopies attenuate the negative drought impact on decomposition due to indirect canopy effects and direct litter mixing effects.

## 2 MATERIALS AND METHODS

### 2.1 Study sites and experimental design

We have chosen four different sites along a latitudinal gradient with contrasted climatic conditions in the south-west part of the French Alps (Fig. S1). The forests at the two northern sites were dominated by European beech and silver fir, and those at the two southern sites were dominated by European beech and downy oak. According to the French National Inventory, beech represents 12% of all monospecific forest stands, and contributes to 38% of all two-species mixed stands and to 92% of all three-species mixed stands across the French Alps. All our study sites have a closed forest canopy and forests grow on limestone bedrock with a north to west exposition, but they vary to some extent in stand and soil characteristics (Table 1).

**Table 1:** Characteristics of the four study sites, including geographic, climatic, forest stand, soil, and soil microclimatic variables. Annual mean temperature and precipitation represent average values during the period of 2015-2018 measured at the nearest meteorological station. Tree height indicates the average height of the dominant trees within the experimental plots and the range (in brackets) among the six plots of each site. Likewise, basal area indicates the average and the range (in brackets) among the six plots per site. Soil characteristics were determined from a bulk sample of four soil cores (to a depth of 15 cm) per plot. Mean values of the six plots per site are shown. Soil microclimate data are from measurements in two plots per site, with a total of four sensors (two per plot) collected during the experiment. Annual mean values represent the average of the two full years covered with our study (2016 and 2017) excluding the rain exclusion subplots for the soil moisture data. Minima and maxima are the average of the four lowest or highest values measured by the four sensors (i.e. including the rain exclusion subplots) at each site during the entire experimental period.

|  | **Study sites** |  |  |  | 717 |
| --- | --- | --- | --- | --- | --- |
|  | **Vercors** | **Ventoux** | **Lubéron** | **Ste Baume** | 718 |
| **Site characteristics** |  |  |  |  |  |
| Latitude, longitude (°) | 44.93°N / 5.32°E | 44.19°N / 5.25°E | 43.82°N / 5.54°E | 43.33°N / 5.77°E |  |
| Slope (°) | 28 | 18 | 30 | 10 | 719 |
| Altitude (m) | 1264 | 1304 | 937 | 748 |  |
| Mean Annual Temp. (°C) | 6.3 | 6.5 | 10.2 | 10.1 |  |
| Mean Annual Prec. (mm) | 1464 | 1216 | 793 | 940 |  |
| **Forest characteristics** |  |  |  |  | 720 |
| Tree height (m) | 25 (17-33) | 18 (15-21) | 15 (13-16) | 26 (21-31) |  |
| Basal area (m<sup>2</sup>/ha) | 52 (40-63) | 57 (53-60) | 37 (33-40) | 52 (40-63) |  |
| **Soil characteristics** |  |  |  |  | 721 |
| Clay content (%) | 31.5 | 54.9 | 47.1 | 39.7 |  |
| Sand content (%) | 24.0 | 9.0 | 12.0 | 26.2 | 722 |
| Soil pH | 5.2 | 6.9 | 7.6 | 7.1 |  |
| Organic matter content (%) | 15.9 | 30.8 | 15.9 | 14.2 |  |
| C/N ratio | 14.3 | 19.9 | 14.6 | 18.3 | 723 |
| **Soil microclimate** |  |  |  |  |  |
| Annual mean temp. (°C) | 6.0 | 7.0 | 10.1 | 10.9 | 724 |
| Minimum temp. (°C) | -0.2 | -0.3 | 0.0 | 1.8 |  |
| Maximum temp. (°C) | 15.8 | 15.8 | 20.7 | 20.5 | 725 |
| Ann. mean moist. (% vol.) | 32.7 | 23.4 | 17.0 | 22.6 |  |
| Minimum moist. (% vol.) | 11.3 | 10.9 | 0.5 | 9.4 |  |
| Maximum moist. (% vol.) | 44.3 | 41.5 | 33.4 | 39.2 |  |

At each of the four sites, there were six 300 m^2^ plots, two of each composed of a monospecific stand of the two dominant local target species (European beech and silver fir at the two northern sites and European beech and downy oak at the two southern sites) and a mixture of the two species (Fig. S1). Within each plot we randomly (but avoiding unsuitable microsites, for example covered with dead wood or rocky areas) selected four subplots where we installed the litterbags for our decomposition study. Two of these subplots were randomly assigned to a summer drought treatment and the two remaining subplots were used as controls. Within each of the subplots, we placed eight litterbags filled with leaf litter representing the canopy composition of the target species (either one of the two local target species dominating the canopy or a mixture of the two species). This yielded a total of 4 sites x 6 plots (two plots of each of the two monospecific canopies and the mixed canopy) x 4 subplots x 8 litterbags = 768 litterbags. To evaluate the contribution of direct litter mixture effects to the overall canopy effects on decomposition we additionally included two litterbags of single species litter of each of the two tree species present at the respective sites in all of the mixture’s plots. This added an additional 128 litterbags (4 sites x 2 plots x 4 subplots x 2 litter types x 2 litterbags) for the mixed species plots. In total, we placed 896 litterbags (224 at each site) in autumn 2015.

### 2.2 Application of extended summer drought

We applied a complete rainfall exclusion to half of the subplots within each plot, during the summer (roughly between end of June and end of September, i.e. for about three months, Fig. S1) using a removable custom-made rainout shelter (see photo in Fig. S1). The removable shelter was constructed with transparent plastic sheets covering an about twice as large area than that used for the placement of litterbags on the forest floor. Upslope of each subplot subjected to summer rainfall exclusion, we fixed the plastic sheets down to the forest floor where we additionally dug a small 10 cm deep ditch to direct runoff from the plastic cover and potentially from the forest floor away from the area with the litterbags. The other three sides were kept open with the plastic sheet 50 to 80 cm above the forest floor to allow unhindered air circulation in order to minimize microclimate effects other than rain exclusion. We continuously monitored soil moisture and temperature at 5 cm soil depth in one of each of the rainfall exclusion and the control subplots in each of the tree species mixture plots (i.e. two plots at each site) using automated sensors (RT-1 and EC-5 or GS-1 sensors for temperature and moisture (TDR sensors), respectively, Decagon Devices Inc., Pullman, WA, USA) storing average values every three hours for the whole duration of the experiment. Due to wild boar damage of the sensors and/or data loggers at the Ste Baume site in the first fall after litterbag installation, there were no microclimate data over winter at this site until we replaced the material in the following spring in 2016.

### 2.3 Litterbag construction

The litterbags were 18cm × 18cm large with a relatively small mesh size on the soil-facing bottom (0.5 × 0.5 mm) and a wider mesh size (5 × 8 mm) on the top. We have chosen a coarser mesh size on top of the litterbags to allow access for the whole decomposer community, including soil macrofauna, and a smaller mesh size on the bottom to avoid gravitational loss of small litter particles. The same type of litterbags was used successfully in a previous study (Joly et al. 2017). Each litterbag was filled with 10 g of air-dried litter material with an equal contribution of each of the two species in mixed species litterbags (i.e. 5 g each). We decided to use a common pool of leaf litter material from the identical species present in our study but collected at different sites than the ones included in our study in order to standardize leaf litter quality across all plots and sites. We did this to avoid confounding between site and plot effects with site-specific litter quality effects. Beech and fir litter were collected in a Romanian forest nearby Râsca (47°36’N, 26°23’E) and oak leaf litter was collected in a Mediterranean forest nearby Montpellier (47°60’N, 3°60’E). All leaf litter was collected with suspended litter traps emptied every two weeks from October 2011 to November 2012 in Romania and from November 2014 to April 2015 in Montpellier.

### 2.4 Litterbag harvest and laboratory analysis

The experiment was set up during late October and early November 2015 and lasted two and a half years when the last litterbags were harvested in May 2018, i.e. including three winters (2015-2016, 2016-2017 and 2017-2018) and two summers (2016 and 2017). We harvested subsets of litterbags five times during the experiment to follow the dynamics of litter C and N loss. We collected one to two litterbags from each subplot at all sites after about 180 days (18^th^ April to 5^th^ May 2016), 360 days (26^th^ September to 20^th^ October 2016), 560 days (4^th^ to 16^th^ May 2017), 700 days (15^th^ to 18^th^ September 2017), and 910 days (23^th^ to 24^th^ April 2018). The respective harvesting periods differed somewhat in duration because of logistics and accessibility to sites, which were far from each other (about 300 km between the most northern and most southern site). The single species litterbags exposed in the mixture plots for the assessment of litter mixing effects were removed from the field during the fourth harvest after 700 days of exposure (only one harvest). All litter and soil material that adhered at the outside of litterbags were carefully removed in the field, each litterbag was then placed in an individual paper bag in the field and air-dried for at least 48 hours using an oven (set at a maximum temperature of 30°C) once back in the lab. The litter was then sorted to species (in mixed litterbags), cleaned from extraneous particles (e.g. pieces of seeds, bud scales), and weighed at a precision of ± 0.01g (Precisa 3610CD, Balco, St Mathieu de Treviers, France). We assessed decomposition by quantifying mass loss relative to initial mass, and C and N loss relative to the initial amount of these elements. For the determination of C and N concentrations in initial and decomposed litter material, we ground all samples to a particle size of < 1mm (FOSS cyclotec mill, ThermoFisher scientific, Waltham, USA), and analyzed the powder with an elemental analyzer (Flash 1112 series EA, ThermoFinnigan, Milan, Italy). We then calculated litter C and N loss as a percentage of the initial amount. Besides C and N concentration, the initial litter material was also analyzed for cellulose, hemicellulose, and lignin determined by sequential extraction according to the Soest and Wine (1967) protocol using a fiber analyzer (Fibersac 24; Ankom, Macedon, NJ, USA), condensed tannins using the acid butanol method (Porter et al. 1986), and the minerals Ca, Mg, Mn, and Fe after mineralization (Ethos One, Milestone, Brøndby, Danemark) using an atomic absorption spectrometer (ice 3000 series AA spectrometer, ThermoFisher scientific, Waltham, USA). Initial litter quality variables for all three species are shown in Table 2.

**Table 2:** Initial litter chemistry of the common litter pool used for the construction of litterbags for the three studied tree species (mean values ± standard deviation). Carbon, nitrogen, cellulose, hemicellulose, lignin, and condensed tannin values are given as percent of total litter dry mass. Ca, Mg, Mn and Fe are given in per mil of total litter mass.

|  | Carbon | Nitrogen | Cellulose | Hemicellulose | Lignin | Condensed tannins | Calcium | Magnesium | Manganese | Iron |
| --- | --- | --- | --- | --- | --- | --- | --- | --- | --- | --- |
| **Fir** | 48.5 ± 0.35 | 1.45 ± 0.22 | 14± 0.6 | 44 ± 5 | 18 ± 1.8 | 3.16 ± 0.33 | 15.6 ± 0.64 | 1.64 ± 0.05 | 1.00 ± 0.06 | 0.45 ± 0.10 |
| **Beech** | 46.9 ± 0.11 | 0.85 ± 0.09 | 18 ± 0.7 | 39 ± 2 | 21 ± 0.4 | 8.07 ± 1.10 | 11.4 ± 1.81 | 1.99 ± 0.05 | 0.41 ± 0.07 | 0.22 ± 0.06 |
| **Pubescent oak** | 43.9 ± 2.91 | 0.85 ± 0.03 | 16 ± 0.6 | 50 ± 2 | 19 ± 0.7 | 3.08 ± 0.71 | 11.5 ± 5.26 | 2.26 ± 0.37 | 0.06 ± 0.01 | 0.47 ± 0.03 |

### 2.5 Data analyses

#### 2.5.1 Mass loss dynamics

Remaining litter mass at the five different harvests over the total of 30 months of field exposure was fitted with a negative single exponential decay model (Olson 1963), which fitted our data better than the other frequently used models, such as the linear decay or the asymptotic decay model (Berg and Laskowski 2005):

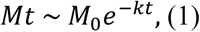

where M_t_ is the mass remaining at time *t*, M_0_ is the litter mass at the beginning of decomposition, and k is the decomposition constant (rate constant).

We analyzed *k* as the response variable with linear mixed-effects models using the “stats” package of the R software (version 3.4.4). We evaluated how much of the variability in *k* was explained by site (*site*), tree canopy composition (*composition*), and rainfall exclusion (*treatment*) according to the model:

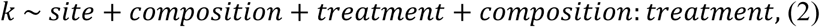

Because the tree species composition, and thus litter identity differed between the northern and the southern part of the gradient we additionally analyzed the two parts of the gradient separately.

#### 2.5.2 Carbon and nitrogen loss

In addition to the decomposition constant for total litter mass loss, we also evaluated C and N dynamics by calculating the amount of C and N remaining at each individual harvest.

Remaining C and N were expressed as a percentage of initial C and N. We analyzed both response variables (C remaining, and N remaining) with linear mixed-effects models using the “stats” package of the R software (version 3.4.4). The fitted explicative variables included site, tree canopy composition, rainfall exclusion, and time (for the five successive harvests).

The full model for C and N loss was:

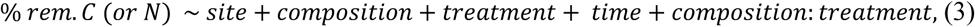

Because the tree species composition, and thus litter identity differed between the northern and the southern part of the gradient we also analyzed the two parts of the gradient separately.

#### 2.5.3 Litter mixture effect vs stand diversity effect

The separation of species-specific leaf litter from litterbags decomposing in the mixed species forest stands and the additional single species litterbags exposed in the same stands, allowed us to calculate the net biodiversity effect (*NBE*) on carbon and nitrogen loss (hereafter “*NBEc*”, “*NBEn*”) after approximately 700 days of litter decomposition (one harvest of single species litterbags only). The net biodiversity effects on total mass and carbon loss were almost identical, and we evaluated only carbon and nitrogen loss in further detail. The *NBE* introduced by Loreau and Hector (2001) is a commonly used metric to quantify biodiversity effects. It compares the observed process rate in mixtures with that predicted from the respective single species treatments by considering the relative abundance of contributing species. We used remaining litter mass as the response variable, therefore, positive *NBE* values indicate antagonistic mixture effects (higher amounts of remaining litter that equals slower decomposition), negative values indicate antagonistic effects, and *NBE* values that are not significantly different from zero mean that there are no mixture effects on remaining litter mass. These *NBE* effects were calculated at the level of individual subplots based on the three litterbag types (mixed leaf litter, and the two mono-specific litterbags) collected during the fourth harvest. In addition, we calculated *NBE* also at the stand level using the mono-specific litterbags from the plots with a mono-specific canopy. This allowed us to compare *NBE* resulting from only litter type interactions within the exact same micro-environment underneath the same mixed-species canopy with *NBE* resulting from all combined effects of mono-specific tree canopies versus mixed tree species canopies, including indirect effects on microclimatic conditions or on decomposer communities.

Differences between *NBE* values of different treatments were evaluated using two-tailed student *t*-tests.

## 3 RESULTS

### 3.1 Rain exclusion during summer

The rainout shelters we installed during the two summers in 2016 and 2017 to simulate more severe and longer summer droughts, appeared to have excluded rainfall efficiently according to the soil moisture measurements (Fig. S2). Additionally, when we were removing the rainout shelters in fall, the litter layer and soil surface were completely dry in all exclusion subplots. In contrast, in control subplots the litter layer and soil surface were often wet when there were recent rainfall events (depending on the site and the year), which was typically the case for example at the Vercors site (Fig. S2). Soil moisture varied strongly over time with clear seasonal differences at all sites (Fig. S2). It was generally lower in the rain exclusion treatments when the rainout shelters were in place during summer and early fall compared to the control treatments across all sites (Fig. S2). However, these differences varied among sites and year depending on the specific rainfall patterns at the different sites and in the two years. For example, the differences were particularly large in the Vercors with more than twice as high mean soil moisture in the control (46.8% and 64.9% of maximum soil water content in 2016 and 2017, respectively) than the exclusion treatment (20.6% and 29.9% of maximum soil water content in 2016 and 2017, respectively) during the period of rainfall exclusion (Fig. S2) because of regular rainfall during the summer. The differences in the other three sites with less frequent summer rain were less pronounced, in particular at the Ventoux and the Lubéron sites with a relatively weak difference of 4 to 5% between the control and the rain exclusion treatment. Soil moisture during the summer was generally low at the Lubéron site in both treatments, ranging between 14.9% in the exclusion subplots in 2017 to 21.5% in the control subplots in 2016 (Fig. S2).

### 3.2 Litter mass loss rates

Litter decomposition proceeded slowly in general with an average of more than 50% of the total initial mass remaining after 30 months of exposure in the field across all sites and for all litter types (Fig. 1). The decomposition constant *k*, determined with the single negative exponential model, varied between 0.08 and 0.26 among sites and litter types (Table S1), and was generally lower when rainfall was excluded (mean of 0.13 across all sites and litter types) compared to control conditions (mean of 0.17 across all sites and litter types). However, despite overall smaller *k*-values in exclusion subplots, there was neither a significant rainfall exclusion, nor a site or canopy composition effect on *k* in the northern part of the gradient (Table 3). In contrast, in the southern part of the gradient rainfall exclusion had an overall negative effect on *k* (Table 3). Additionally, *k* differed significantly between the two sites with higher *k* at Ste Baume compared to Lubéron. Moreover, monospecific beech stands had lower *k* than monospecific oak stands or mixed species stands, but the latter two did not differ.

**Table 3:** Effects of *site* (two for each part of the gradient), tree canopy composition (*composition*), with beech and the second species (other, i.e. fir in northern part and oak in southern part) compared to the mixture, and rain exclusion (*treatment*) on the decomposition constant (k). Coefficient estimate, with standard deviation (in brackets), and part of variance explained are shown. The northern (sites Vercors and Ventoux) and southern (sites Luberon and Ste-Baume) part of the gradient are analyzed separately. Significant effects (p < 0.05) are shown in bold.

|  |  | **northern part** |  | **southern part** |  |
| --- | --- | --- | --- | --- | --- |
|  |  | **Estimate** | **% variance explained** | **Estimate** | **% variance explained** |
| **Intercept** |  | **0.12 (±0.03)** | **26** | **0.24 (±0.02)** | **67** |
| **Site** |  | 0.02 (±0.02) | - | **-0.09 (±0.01)** | **15** |
| **Composition** |  |  |  |  |  |
|  | **Beech** | 0.05 (±0.04) | - | **-0.05 (±0.02)** | **3** |
|  | **Other** | 0.02 (±0.04) | - | 0.02 (±0.02) | - |
| **Treatment** |  | -0.03 (±0.04) | - | **-0.05 (±0.02)** | **2** |
| **Composition : Treatment** |  |  |  |  |  |
|  | **Beech** | -0.03 (±0.05) | - | 0.01 (±0.03) | - |
|  | **Other** | -0.06 (±0.05) | - | 0.04 (±0.03) | - |

**Figure 1:**
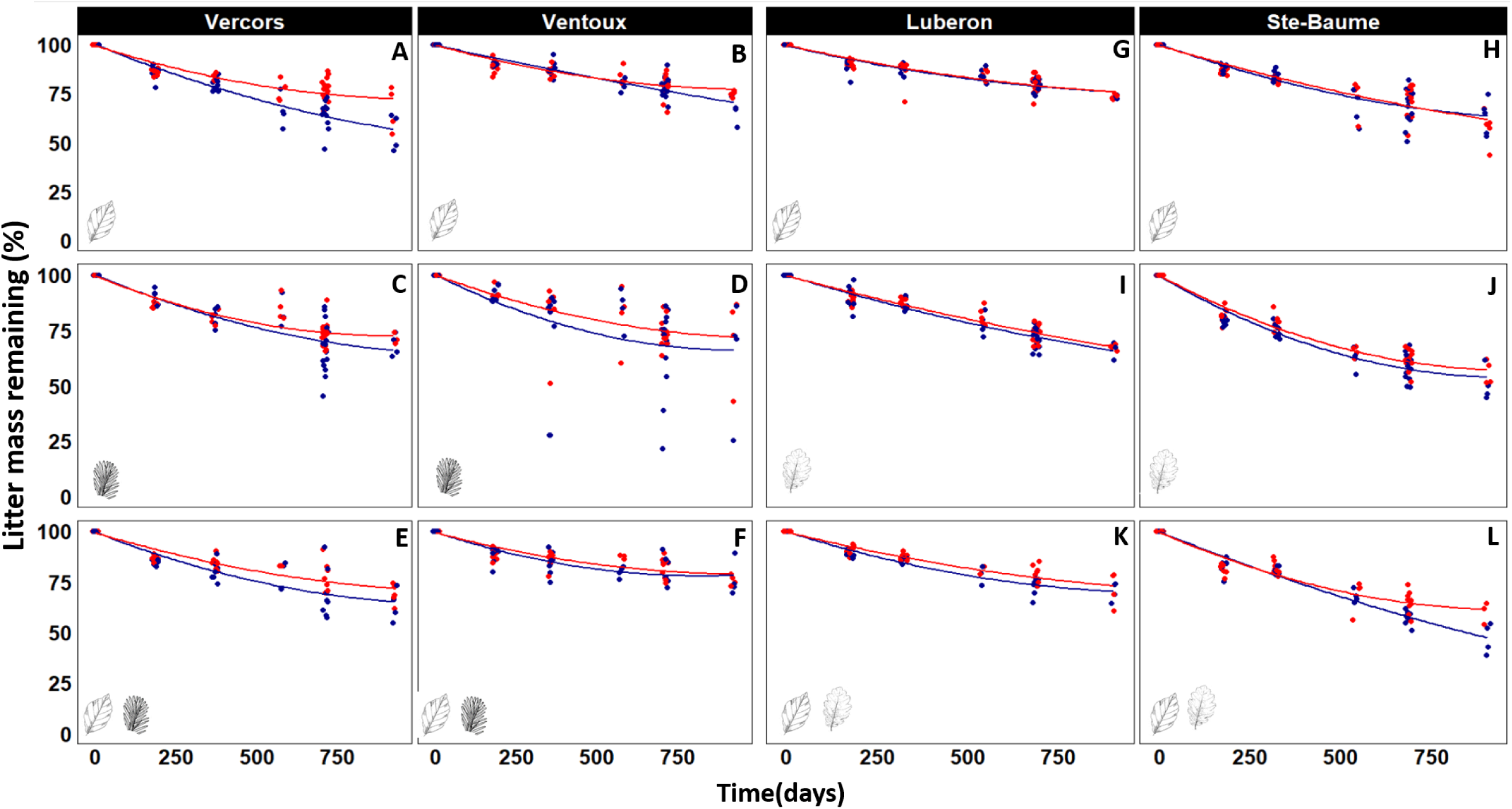
Total litter mass remaining (%) as a function of time (days) for the northern (left panels A-F) and the southern (right panels G-M) part of the gradient for each site and each tree species composition (indicated with the respective leaf symbols in the bottom left corner of the graphs). Blue symbols and lines denote the control and red symbols and lines the rainfall exclusion treatment. Lines represent the average negative single exponential fit of each site, stand composition and treatment.

Since beech was present along the whole gradient, we additionally analyzed the effects of site, canopy composition (pure beech vs. mixture), and rainfall exclusion on beech leaf litter decomposition across the whole gradient. We found significantly slower beech leaf litter decomposition at the Ventoux and the Lubéron site compared to the Vercors and the Ste Baume site (Fig. S3, Table S2). Rainfall exclusion had an overall significant negative effect on *k*, and the canopy composition influenced beech leaf litter decomposition depending on the associated species (Table S2). In mixtures with fir *k* was lower compared to pure beech stands, but there was no difference between pure beech stands and mixtures with oak.

### 3.3 Carbon and nitrogen dynamics

Initial leaf litter C concentration varied somewhat among leaf litter of the three species (Table 2), with decreasing [C] in the order of fir, beech and then oak, with however, no significant difference (tested with Student test) between beech and oak. Similarly, initial leaf litter N concentration also was highest in fir with no significant differences (tested with Student test) between beech and oak. In addition to N, fir litter tended to be richer in nutrient concentrations compared to the two broadleaf species, particularly for calcium and manganese (Table 2). Differences among species were less pronounced for magnesium and iron. Beech showed the highest concentrations of cellulose, condensed tannins and lignin compared to the other two species, and oak had the highest concentration of hemicellulose of all three species.

Carbon dynamics during decomposition proceeded largely in parallel to mass loss and with the same patterns as those observed for litter mass loss rates. Across all sites and litter types, an average of 35% C was lost after 30 months of decomposition. Due to the steadily decreasing quantity of C with ongoing decomposition, the factor “time” had the strongest effect and accounted for the largest part of variance in both parts of the gradient (Table 4 and Fig. 2). Sites differed in C loss dynamics, with a strong difference in the southern part of the gradient (Table 4) where less C was lost from decomposing litter at the Lubéron site (Fig. 2). Rain exclusion led to lower C loss at all sites, but this negative effect was more pronounced in the northern part of the gradient (Table 3). Litter types differed in C loss dynamics in the southern part but not in the northern part of the gradient. Specifically, in pure beech stands in the South, less C was lost from decomposing leaf litter compared to pure oak stands or mixed beech-oak stands, with the latter two showing similar C loss dynamics. The selected best models for C loss dynamics have r-squared of 0.48 and 0.82 in northern and southern part, respectively (Table 4).

**Table 4:** Effects of *site* (two for each part of the gradient), tree canopy composition (*composition*), with beech and the second species (other, i.e. fir in northern part and oak in southern part) compared to the mixture, rain exclusion (*treatment*), and time of harvest (*time*) on total remaining carbon. Coefficient estimate, with standard deviation (in brackets), and part of variance explained are shown. The northern (sites Vercors and Ventoux) and southern (sites Luberon and Ste-Baume) part of the gradient are analyzed separately. Models r-squared are 48% and 81% for northern and southern part, respectively. Significant effects (p< 0.05) are shown in bold.

|  | northern part |  | southern part |  |
| --- | --- | --- | --- | --- |
|  | Estimate | % variance explained | Estimate | % variance explained |
| **Intercept** | **0.92 ($\pm 0.02$)** | **84** | **0.86 ($\pm 0.01$)** | **81** |
| **Site** | **-0.04 ($\pm 0.01$)** | **1** | **0.09 ($\pm 0.01$)** | **3** |
| **Composition** |  |  |  |  |
| **Beech** | -0.01 ($\pm 0.02$) | - | **0.04 ($\pm 0.01$)** | **0** |
| **Other** | 0.01 ($\pm 0.02$) | - | -0.01 ($\pm 0.01$) | - |
| **Treatment** | **0.05 ($\pm 0.02$)** | **0** | **0.03 ($\pm 0.01$)** | **0** |
| **Time (days)** | **0.0003 ($\pm 0$)** | **7** | **0.0004 ($\pm 0$)** | **12** |
| **Composition : Treatment** |  |  |  |  |
| **Beech** | 0.0001 ($\pm 0.03$) | - | -0.02 ($\pm 0.02$) | - |
| **Other** | -0.01 ($\pm 0.03$) | - | -0.02 ($\pm 0.02$) | - |

**Figure 2:**
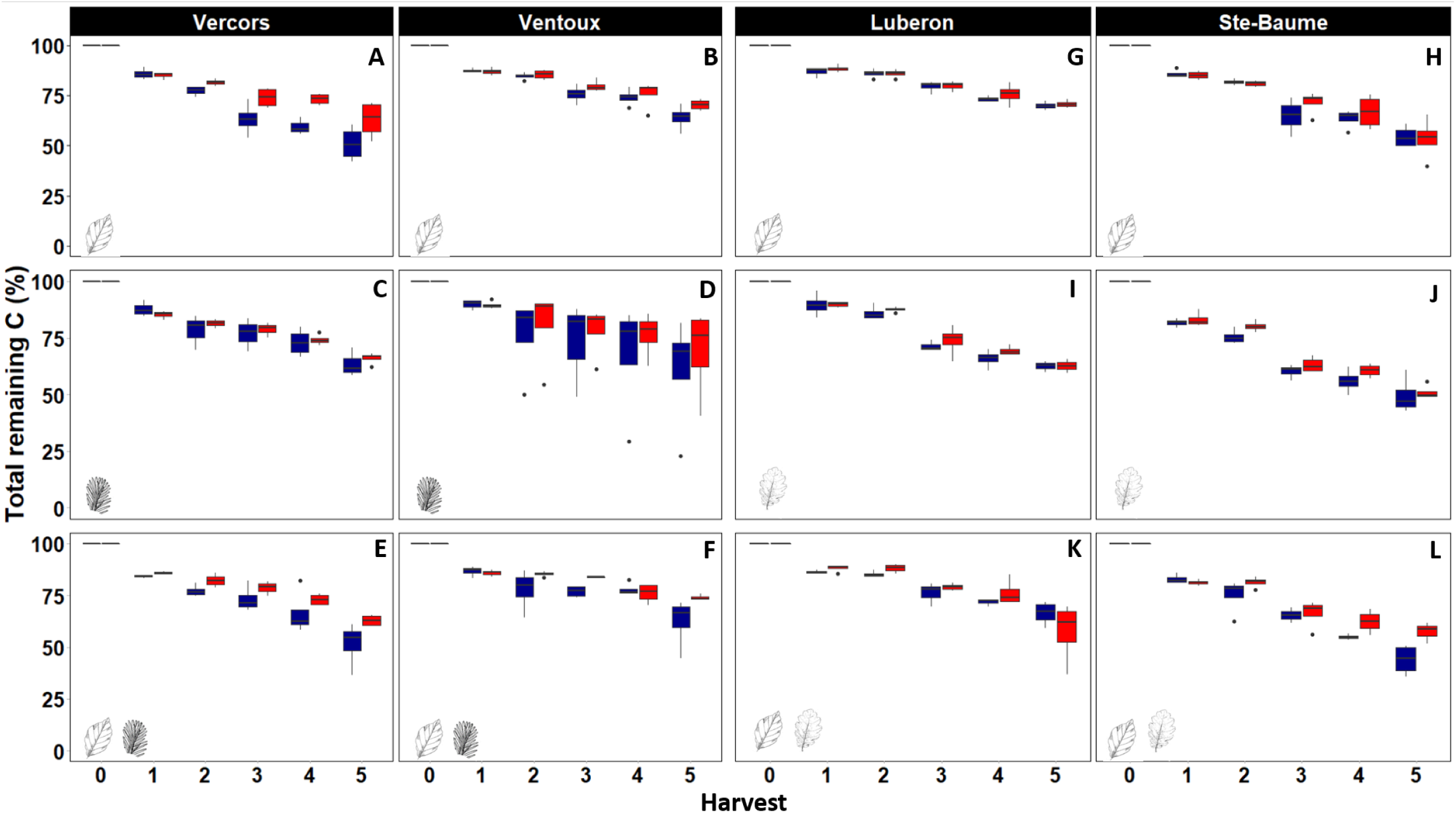
Total C remaining for the five consecutive harvests (i.e. 1 and 2 for Spring and Fall 2016, 3 and 4 for Spring and Fall 2017 and 5 for spring 2018) for the northern (left panels A-F) and the southern (right panels G-M) part of the gradient for each site and each tree species composition (indicated with the respective leaf symbols in the bottom left corner of the graphs). Boxplots in blue denote the control and those in red the rain exclusion treatment.

Nitrogen dynamics differed considerably from C dynamics with overall low N losses of an average 15% across all sites and litter types after 30 months of decomposition. Most importantly, total N remaining in decomposing litter varied little over time (Fig. 3) and consequently, the factor “time” accounted for much less variance compared to the models for C dynamics (Table 5). On the other hand, the litter type had a much stronger influence on N compared to C dynamics, especially in the northern part of the gradient where litter type accounted for 10% of the variance compared to 2% in the southern part (Table 5). In both parts of the gradient across all sites we measured the lowest N loss rates from decomposing beech leaf litter in pure beech stands compared to the other litter types. In the northern part of the gradient fir litter lost the highest amount of N and beech-fir mixtures intermediate amounts. Likewise, in the southern part of the gradient, the highest amount of N was lost from pure oak leaf litter, with intermediate loss rates in mixed beech-oak litter. The site effects were comparatively weak but differed significantly with lower N loss at the Lubéron compared to the Ste Baume site (Table 5) and at the Ventoux compared to Vercors. In contrast to C dynamics, there was no rainfall exclusion effect on N dynamics. Overall, the selected best models for N loss dynamics have r-squared of 0.56 and 0.44 in northern and southern part, respectively (Table 5).

**Table 5:**
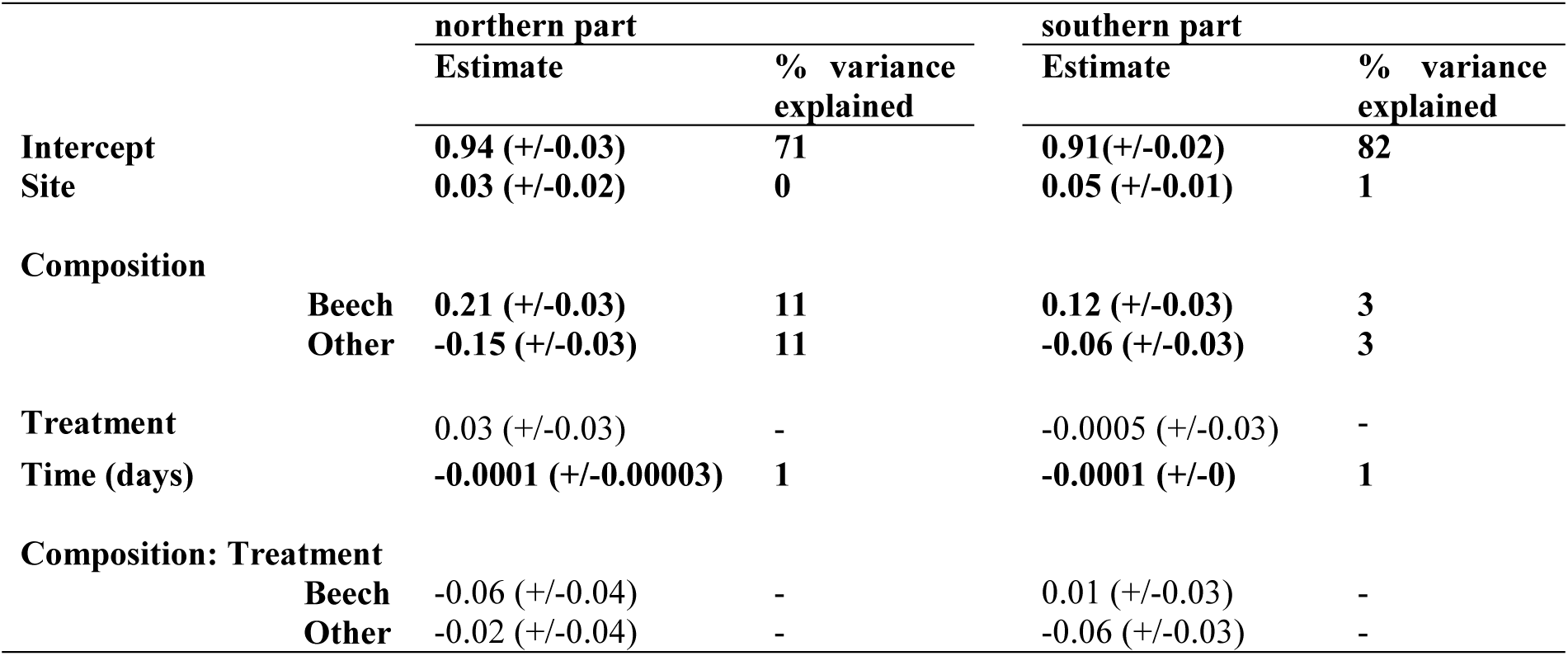
Effects of *site* (two for each part of the gradient), tree canopy composition (*composition*), with beech and the second species (other, i.e. fir in northern part and oak in southern part) compared to the mixture, rain exclusion (*treatment*), and time of harvest (*time*) on total remaining nitrogen. Coefficient estimate, with standard deviation (in brackets), and part of variance explained are shown. The northern (Vercors and Ventoux) and southern (Luberon and Ste Baume) part of the gradient were analyzed separately Models r-squared are 56 % and 44% for northern and southern part, respectively. Significant effects (p < 0.05) are shown in bold.

**Figure 3:**
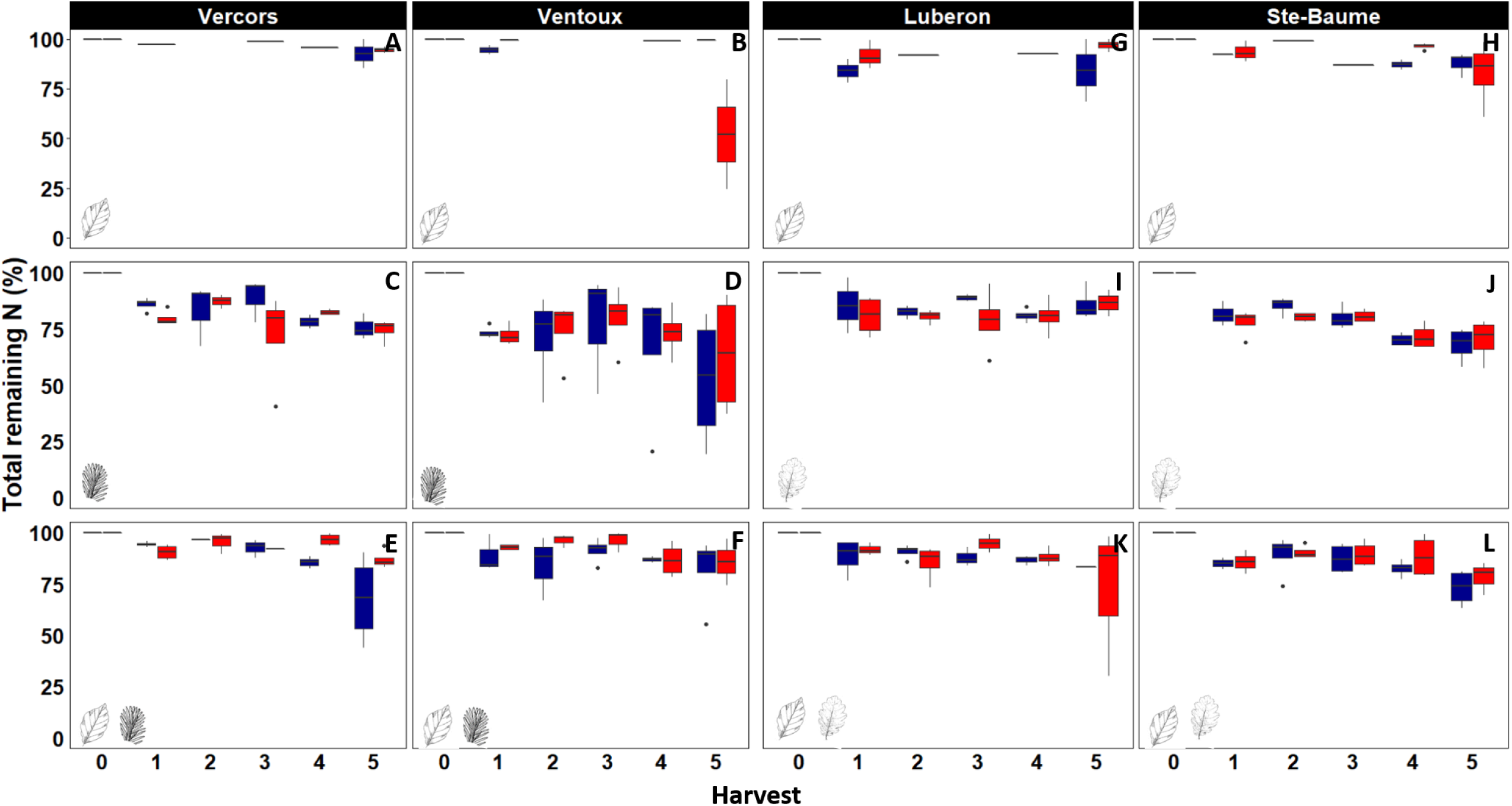
Total N remaining for the five consecutive harvests (i.e 1 and 2 for Spring and Fall 2016, 3 and 4 for Spring and Fall 2017 and 5 for spring 2018) for the northern (left panels A-F) and the southern (right panels G-M) part of the gradient for each site and each tree species composition (indicated with the respective leaf symbols in the bottom left corner of the graphs). Boxplots in blue denote the control and those in red the rain exclusion treatment.

### 3.4 Net biodiversity effects on decomposition

The mixing of litter within the mixed species forest stands under otherwise the same micro-environmental conditions led to antagonistic effects (slower decomposition, i.e. higher amounts of C and N remaining) in mixtures of beech and fir litter (positive *NBE* on remaining C and N, Fig. 4a, 4c), which was mostly driven by fir for C dynamics and by beech for N dynamics. Across forest stands, i.e. when pure beech and pure fir forests are compared to a mixed stand of these two species, there were no *NBE* on remaining C (Fig. 4a), and remaining N (Fig. 4c). The mixture effects differed in the southern part of the gradient. The *NBE* on remaining C (Fig. 4b) and N (Fig. 4d) in beech leaf litter were negative both within mixed species stands and across forest stands, meaning that more C and N was lost from decomposing beech leaf litter in mixtures compared to mono-specific beech litter. Oak leaf litter showed opposite *NBE* compared to beech that is less C and N loss from decomposing oak litter when it was mixed with beech than when it decomposed singly (Fig. 4b, 4d). This antagonistic effect on oak leaf litter decomposition was somewhat more pronounced across stands (i.e. when pure oak litter decomposed underneath a mono-specific oak canopy) compared to that within the mixed tree species stand. Consequently, the synergistic effects on beech C and N loss were largely cancelled out by oak across stands, while it persisted underneath a common mixed tree species canopy (Fig. 4b, 4d).

**Figure 4:**
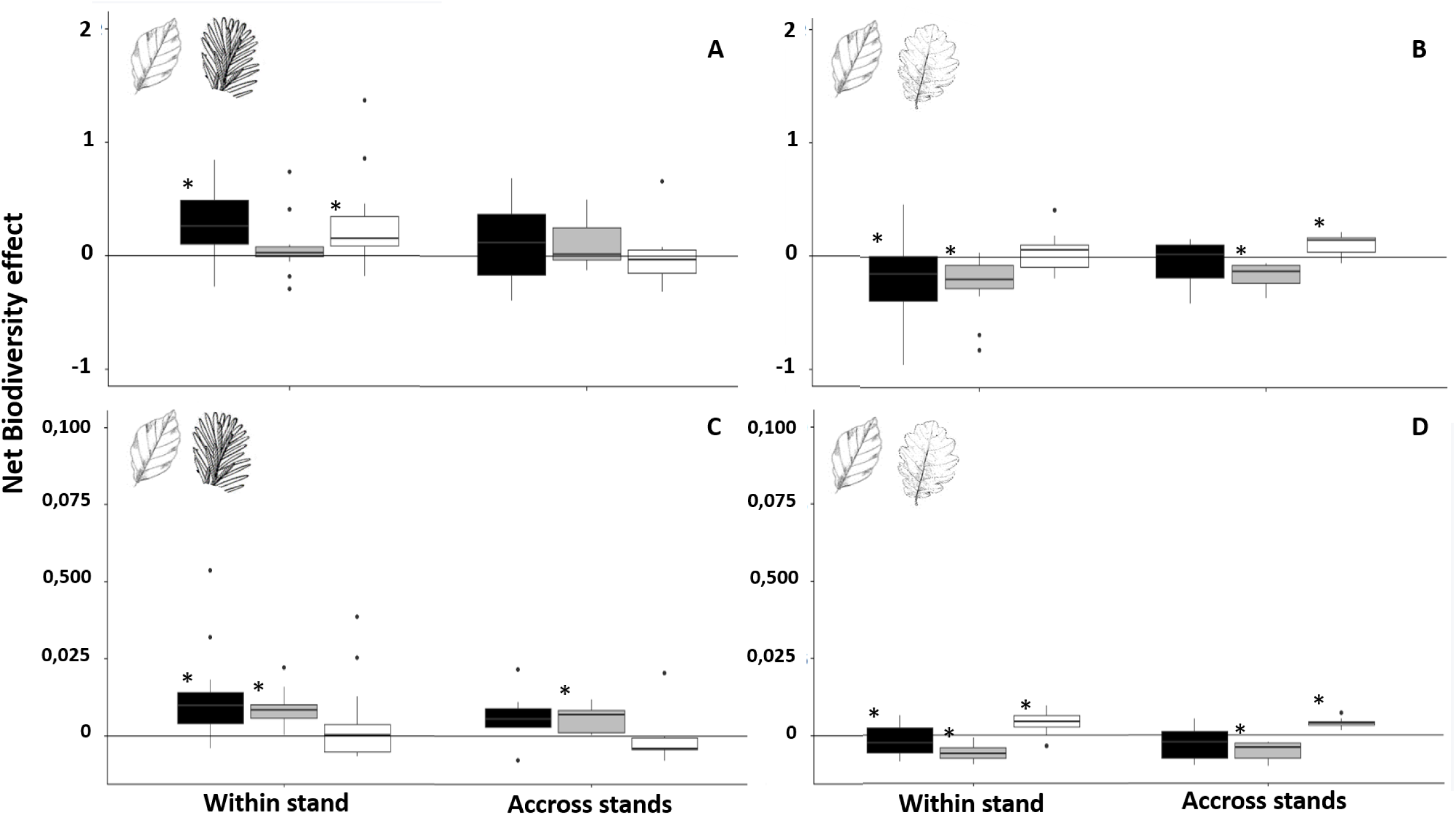
Net biodiversity effects (NBE) on decomposition (note that here it was expressed as remaining mass) within a forest stand (i.e. single litter species and mixed leaf litter decomposing underneath the same mixed tree species canopy) or across forest stands (i.e. single litter species decomposing underneath mono-specific canopies and litter mixtures decomposing underneath mixed tree species canopies). The top panels (A) and (B) show NBE for total C remaining in beech-fir and beech-oak mixtures, respectively. The bottom panels (C) and (D) show NBE for total N remaining in beech-fir and beech-oak mixtures, respectively. Total mixture NBE (black boxes), beech litter NBE (grey boxes), and fir litter NBE (white boxes in panels A, C) or oak litter NBE (white panel in panels B, D) are shown separately. The horizontal line (y=0) indicates no net biodiversity effect, and positive or negative values show positive NBE (antagonistic, i.e. mixture slows decomposition) or negative NBE (synergistic, i.e. mixture accelerates decomposition), respectively (the counterintuitive sign of the diversity effect results from the response variable that is remaining C or N). Stars (*) indicate NBE that are significantly different from zero (p < 0.05, Student test).

## 4 DISCUSSION

### 4.1 Effects of a longer and more severe summer drought

We hypothesized a negative effect of longer and more severe summer droughts on leaf litter decomposition (i.e. decreased mass loss rates and less carbon and nitrogen release), as soil moisture has a strong influence on decomposer organisms, and thus, on decomposition (Aerts 1997). Our hypothesis is partially supported by the data we collected across four different sites covering a north-south gradient in the Western French Alps. Overall, the total exclusion of rainfall during approximately three months in two consecutive summers (i.e. 6 months out of a total 30 months of decomposition) had a relatively small effect on decomposition. The decomposition rate constants calculated from five harvests of litterbags were lower with rainfall exclusion in the southern part of the gradient but accounted for an only small amount of total variation. Opposite to our prediction, *k* was not significantly different under rainfall exclusion compared to controls in the northern part of the gradient. The response to summer drought was more consistent for carbon loss, with a negative effect in both parts of the gradient, but still accounting for only a small amount of the overall variation. In contrast, nitrogen loss did not change in response to an extended summer drought irrespective of the location of the study sites.

A drier climate had a direct negative effect on litter mass loss and C release in previous studies in different types of ecosystems and using different approaches of precipitation manipulations, for example in a temperate forest using irrigation (Lensing and Wise 2007), in Mediterranean forests and shrublands applying total rainfall exclusion (Saura-Mas and others 2012; Santonja and others 2017, 2019), in a temperate grassland also with total rain exclusion during a part of the year (Vogel and others 2013), or in a tropical forest with partial exclusion (Wieder and others 2009). There may be different reasons for why we did not observe stronger effects of extended summer drought. In a field experiment like ours, the amount of excluded rainfall, and thus its potential impact on decomposition, depends strongly on the frequency and amount of precipitation during the rain exclusion period. Indeed, at some of our sites like Lubéron and Ventoux there was very little precipitation during the summer and early fall months when we applied our rain exclusion treatment, which was seen in the soil moisture measurements with comparatively small differences between the two treatments (Fig. S2). Contrariwise, at the Vercors and Ste Baume sites, there were more regular rainfall events during the exclusion period resulting in clearly higher soil moisture in the control treatment (Fig. S2). This suggests that the three months of total rain exclusion we applied, reinforced summer drought relatively weakly in some of our study sites and consequently limited drought effects on decomposition. The differences in summer rainfall patterns were not neatly distinguished between the northern and the southern part of the gradient, which is probably the main reason why extended summer drought had not a stronger effect in the northern than the southern part as we initially hypothesized.

The proportion of litter mass loss during summer relative to the whole year may be small generally, with most of the decomposer activity in spring and late fall. Seasonal decomposer activity was shown in other studies from temperate forests (Bradford and others 2008; Kang and others 2009) and even more so in Mediterranean forests (Hornung and Warburg 1995; Fioretto and others 2001; Santonja and others 2017). Indeed, the relative changes in litter mass under control conditions were overall larger between the late fall and spring collections of litterbags than in the other half of the year (Fig. 2, Fig. S3), especially for beech with an average of 14% change between late fall and spring compared to 4% between spring and late fall. Such intra-annual differences in decomposition then determine the potential of the effect a precipitation exclusion can maximally have depending on when exactly rainfall is excluded. Changes in precipitation in other parts of the year may have greater consequences for decomposition than what we simulated here. This highlights a critical aspect of rain manipulation studies that may not be sufficiently acknowledged. The timing and type of exclusion (e.g. partial rain exclusion all year round vs. total exclusion only part of the year) can have a strong impact on the effect size, which may complicate direct comparisons among studies manipulating precipitation differently. For example, modifying both, the quantity and frequency of rainfall in a desert ecosystem, Joly and others (2017a) reported that the distribution of rainfall events is as important for decomposition as the absolute amount of rainfall, with different thresholds for distinct groups of decomposer organisms (Joly and others 2019). The consequences of changing precipitation on decomposition therefore may be strongly dependent on the type of ecosystem and the specific predictions for future scenarios under climate change. In the geographical area we studied, more extreme drought periods during the summer seem the most likely modifications in precipitation patterns in the future (Giorgi and Lionello 2008; Dubrovský and others 2014; Polade and others 2017), although precipitation is notoriously difficult to model (Xuan and others 2009). For a better understanding and prediction of climate change impacts on decomposition, it would then make little sense to reduce precipitation with a partial rain exclusion over the whole year or excluding rainfall in autumn.

An inherent characteristic of decomposition studies using litterbags in the field is the large variability among litterbags, which typically is unaccounted for by any treatment factors. Most of this variability likely reflects environmental heterogeneity at the very small spatial scale of individual litterbags (Bradford and others 2016). For example, in our study, the forest floor at the Ventoux site was densely covered with loose gravel from the eroding limestone bedrock. In combination with strong wild boar activity, parts of the subplots were sometimes covered with a dense layer of gravel (which we removed each time we visited the site) modifying micro-environmental conditions for some of the litterbags. Such disturbance may also have led to some litter particle loss. This might have been an issue with fir needle litter in particular, explaining the overall larger variability among litterbags at this site. The considerable spread of the data within a site and a treatment, then makes it more difficult to detect effects at a manageable level of replication with multiple sites. Interestingly, when we ran the statistical analysis across the whole gradient for beech leaf litter decomposition only, the drought effect was significantly negative and accounted for the same amount of variance as site differences and canopy tree species composition (Fig. S3, Table S2). This suggests, that despite the numerous other factors influencing litter decomposition, longer periods of summer drought predicted for the near future, will slow down leaf litter decomposition in the studied forest ecosystems.

Lower decomposition rates with more severe summer drought may have stronger impacts on the C cycle than on N release from decomposing litter, because in contrast to C loss, N loss was not affected in our study. Nitrogen dynamics differed fundamentally from C loss dynamics in our study, as is generally the case and well documented in the literature (e.g. (Kalbitz and others 2000; Berg and Laskowski 2005). In fact, we observed little net N loss with an average of 15% across all sites and forest stands during the experiment. This comparatively low amount of N release may be explained by an extended period of apparent N immobilization in the beginning of the decomposition process (Fig. 3). There were only small differences in N remaining among the different harvests, especially in the first half of the experiment, and net N immobilization was particularly marked in beech leaf litter. Based on our data it seems that C and N dynamics may be more unbalanced with increasing drought as it was shown in previous studies (Alessandro and Nyman 2017; Durán and others 2018), at least during the beginning of the decomposition process, which could have longer term consequences for decomposer communities and biogeochemical cycling at the ecosystem scale. However, studies covering late stages of decomposition are needed for a better understanding of drought effects on N dynamics, when absolute N losses start dominating N dynamics over N immobilization.

### 4.2 Tree mixture effects on decomposition

Our second hypothesis stating that tree species mixtures would attenuate negative drought effects on decomposition was not supported by our data. The main reason for this was probably the weak overall drought effects we observed across the different study sites. On the background of weak drought effects and high unexplained variation in our data, potentially small and subtle indirect canopy composition and direct litter mixing effects on decomposition were difficult to detect. Our hypothesis was based on the few observations of positive diversity effects of mixed plant canopies on the decomposition environment (Hector 1999; Joly and others 2017b), and of mixed litter on microclimatic conditions (Makkonen and others 2013), which explained more rapid decomposition in mixed compared to single species communities previously. There are only very few studies that experimentally manipulated plant diversity and rain exclusion in the same experiment (Santonja and others 2017, 2019) with contrasting results.

Irrespective of the drought treatment, we found significant tree species mixture effects on C and N loss. The direct litter mixing effect determined from litterbags exposed within the same mixed forest stands was generally stronger compared to the overall tree species mixture effect across forest stands composed of either species or a mixture of the two species. When beech litter was mixed with fir litter at the two northern sites, C and N losses were lower, but when it was mixed with oak litter at the two southern sites, C and N losses were higher compared to single litter species. This indicates that direct litter mixing effects depend on mixture composition, which is a common observation in litter mixture studies (e.g. Wardle and others 2003; Barantal and others 2014). Since mixing litter of a broadleaf deciduous species with that of a conifer species often stimulated decomposition of the latter in past studies (see review by Hättenschwiler and others 2005), we expected positive direct litter mixing effects in beech-fir rather than in beech-oak forests. However, C loss from fir litter was slower when it decomposed together with beech compared to when it decomposed alone. This negative mixture effect on fir needle decomposition was no longer observed at the stand level, i.e. when each of the litter type decomposed in its home plot with corresponding canopy composition. In beech-oak forests, C and N loss was stimulated in beech when it decomposed together with oak with opposite effects on oak leaf litter. The net effect on mixture decomposition was still positive though, and the patterns were the same, although weaker, at the stand level when each litter type decomposed in its home plot with corresponding canopy composition. Without additional data on abundance and composition of decomposer organisms, it is difficult to specify the driving mechanisms for opposing diversity effects in beech-spruce and beech-oak forests. Resource complementarity in mixed litter as a key mechanism for litter mixing effects (Hättenschwiler and others 2005; Barantal and others 2014), would have been expected rather in beech-fir mixtures with overall more contrasting litter quality, especially N concentration, between beech and fir than between beech and oak.

Whatever the underlying mechanisms for the observed tree mixture effects on decomposition, its effect size was similar or greater compared to that of severe summer drought. This is an important result, suggesting that climate change driven modifications in tree diversity and composition of forests may have at least as important indirect consequences on decomposition as those in response to direct climate related factors. However, the relative importance of indirect plant community driven and direct climate change effects needs to be interpreted cautiously, because climatic factors are likely more complex than our simulated longer drought period during summer and litter diversity effects may change over time (Santonja and others 2019) and differ if the whole decomposition process is considered.

## AUTHORS’ CONTRIBUTIONS

MJ and SH designed the research and methodology, analyzed the data and wrote the manuscript.

## ACKNOWLEDGMENTS

MJ was supported by an ADEME PhD grant. This study was funded through the project DISTIMACC (ECOFOR-2014-23, French Ministry of Ecology and Sustainable Development, French Ministry of Agriculture and Forest) and the ANR project BioProFor (contract no. 11-PDOC-030-01). We thank D. Degueldre, S. Coq, J. Nahmani, X. Morin, and A. Arriaga for their help during the experimental setup, field work, and sample collections, R. Beugnon, G. Ayache, M. Guers and L. Delteil for assistance in sample preparation and laboratory analyses. All chemical analyses were done at the Plate-Forme d’Analyses Chimiques en Ecologie, LabEx Centre Méditerranéen de l’Environnement et de la Biodiversité, with the help of B. Buatois.

## Notes

### Competing Interest Statement

The authors have declared no competing interest.

