## Supplementary Materials for "Decomposition in mixed beech forests in the south-western Alps under severe summer drought"

Figure S1: Geographical location of the four study sites (a and b) (with blue crosses in fir-beech forests and with red crosses in downy oak-beech forests) of the decomposition study. At each site, we determined two plots of each of the two locally dominant species and their respective mixtures (c), yielding six plots per site. In each individual forest plot, four subplots were installed where litterbags were placed, two of them kept as controls, and two of them equipped with rain shelters during summer - see photographs (d) for an example of a control subplot and (e) for an example of a rain exclusion subplot). The time axis underneath the figures (f) indicates the experimental duration (30 months), the five consecutive harvests (H1-5) and the application of the rainfall exclusion (in orange).

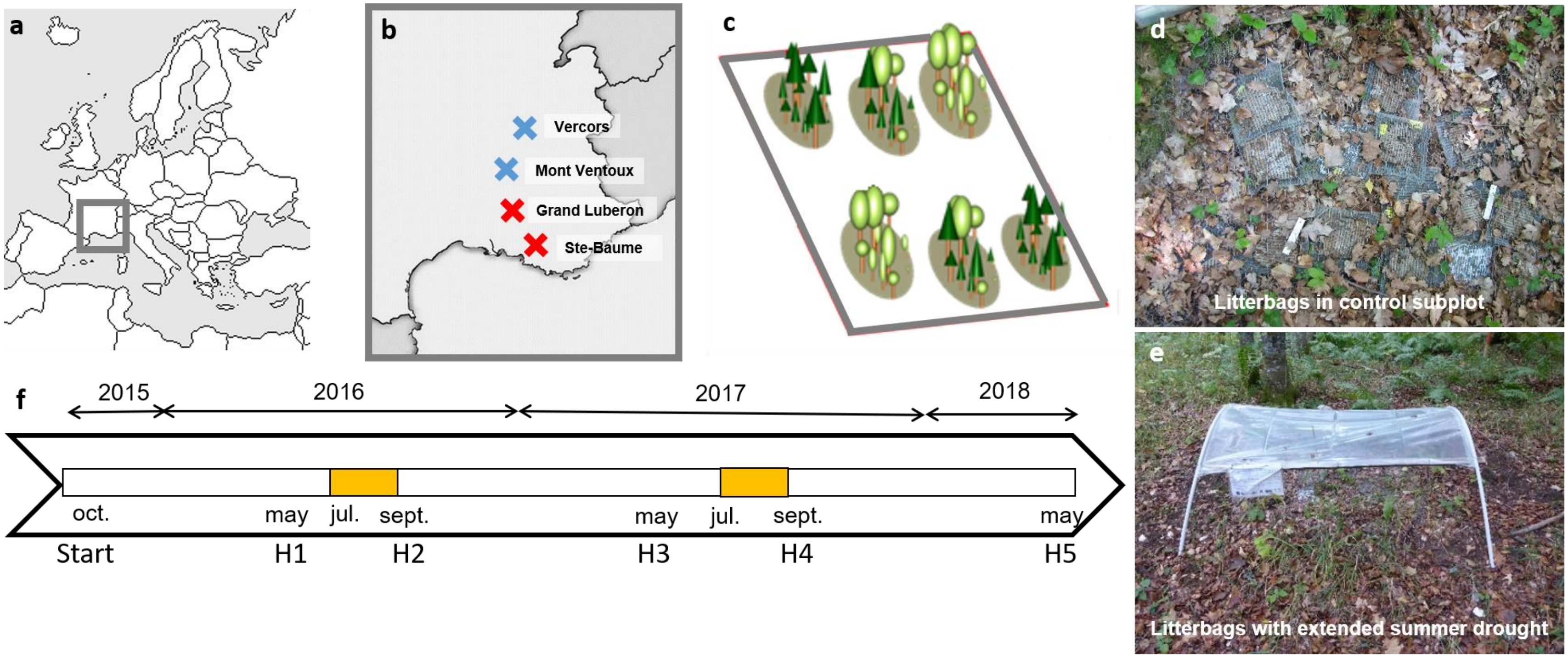

Figure S2. Soil moisture (at 5 cm soil depth) over the entire experimental duration for the four study sites Vercors (A), Ventoux (B), Lubéron (C), and Ste-Baume (D). Daily mean values of two control plots (blue) and two plots with rainfall exclusion (red) per site are shown. The values are in percent of the absolute maximum (10 highest individual measurements taken every three hours during the whole experiment) measured in each individual subplot (i.e. individual sensor specific maximum that is supposed to be at maximum water holding capacity of the respective soil). The fine horizontal black line indicates 40 % of the absolute maximum, a value assumed to impose severe drought stress to soil microorganisms. The grey frames with horizontal grey bars indicate the period with complete rain exclusion in the rainfall exclusion treatment each year (2016 and 2017). For each year the average soil moisture measured in the two treatments from the onset of the rain exclusion until the soil in the exclusion treatment reached 40 % of the absolute maximum after the removal of the rain exclusion, is indicated in each graph. Note that values are missing during the first winter 2015/16 at the Ste-Baume site due to wild boar damage of sensors and data loggers.

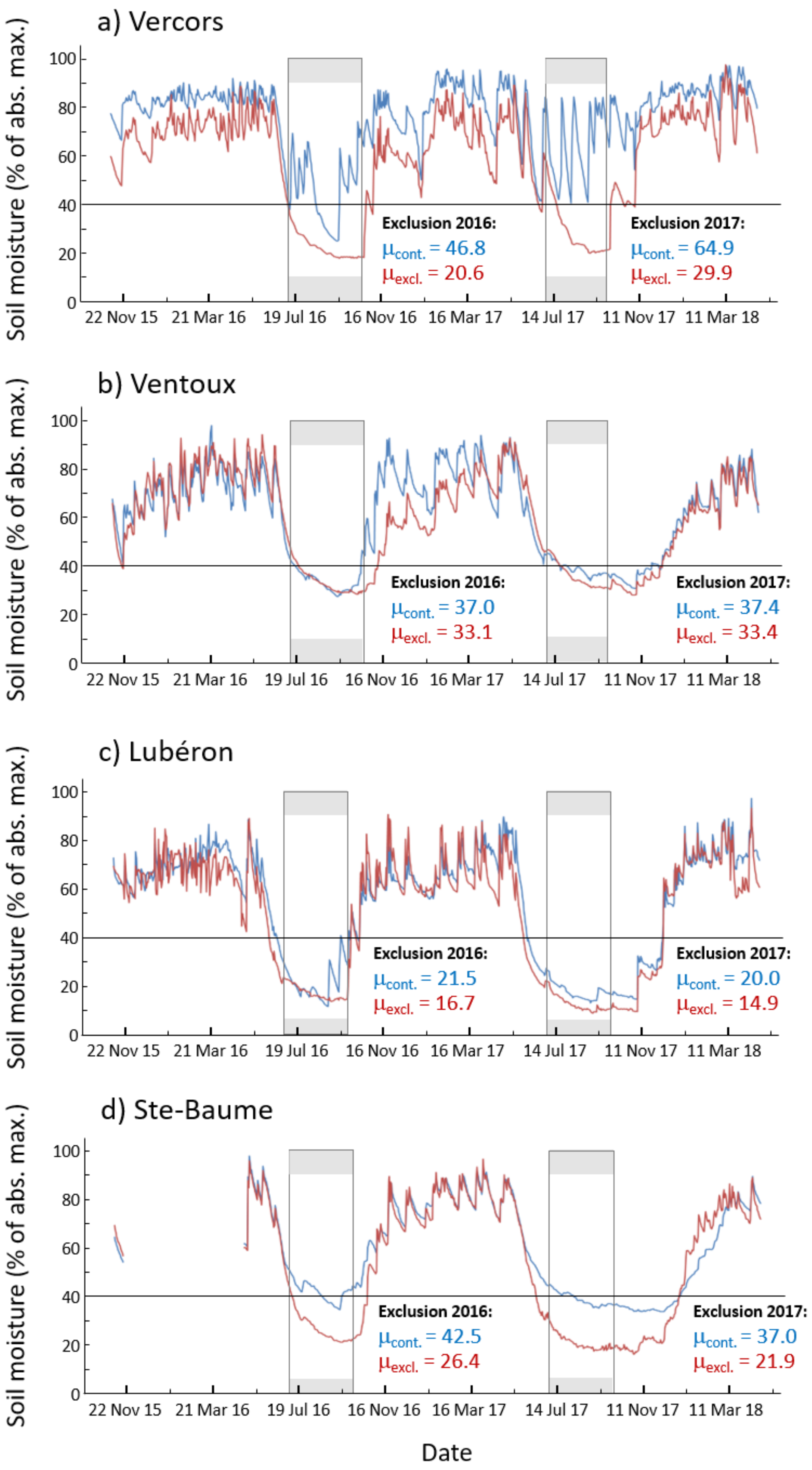

a) Vercors
b) Ventoux
c) Lubéron
d) Ste-Baume
Soil moisture (% of abs. max.)
0
20
40
60
80
100
22 Nov 15
21 Mar 16
19 Jul 16
16 Nov 16
16 Mar 17
14 Jul 17
11 Nov 17
11 Mar 18
Exclusion 2016:
μcont. = 46.8
μexcl. = 20.6
Exclusion 2017:
μcont. = 64.9
μexcl. = 29.9
Exclusion 2016:
μcont. = 37.0
μexcl. = 33.1
Exclusion 2017:
μcont. = 37.4
μexcl. = 33.4
Exclusion 2016:
μcont. = 21.5
μexcl. = 16.7
Exclusion 2017:
μcont. = 20.0
μexcl. = 14.9
Exclusion 2016:
μcont. = 42.5
μexcl. = 26.4
Exclusion 2017:
μcont. = 37.0
μexcl. = 21.9
Date

Table S1. Decomposition rate constants ($k$, $yr^{-1}$) calculated from remaining litter mass determined with five consecutive harvests ($Mt \sim M_0 e^{-kt}$) for each site, treatment and canopy composition separately (mean ± SD).

| Site | Treatment | Beech | Fir | Oak | Mixed |
|---|---|---|---|---|---|
| **Ventoux** | **Control** | 0.14(±0.04) | 0.18(±0.21) | - | 0.09(±0.05) |
| | **Exclusion** | 0.09(±0.03) | 0.11(±0.09) | - | 0.09(±0.03) |
| **Vercors** | **Control** | 0.23(±0.05) | 0.12(±0.03) | - | 0.17(±0.06) |
| | **Exclusion** | 0.11(±0.06) | 0.08(±0.03) | - | 0.12(±0.05) |
| **SteBaume** | **Control** | 0.21(±0.06) | - | 0.25(±0.06) | 0.26(±0.05) |
| | **Exclusion** | 0.18(±0.07) | - | 0.20(±0.04) | 0.17(±0.03) |
| **Luberon** | **Control** | 0.09(±0.02) | - | 0.17(±0.04) | 0.13(±0.02) |
| | **Exclusion** | 0.10(±0.02) | | 0.15(±0.02) | 0.12(±0.03) |

Figure S3. Remaining leaf litter mass of beech at all four sites across the whole gradient (from left to right showing sites from the North to the South). Boxplots in blue denote the control treatment and those in red the rain exclusion treatment. The upper four panels (A, B, E, F) show beech leaf litter mass in the mono-specific beech stands. The lower four panels (C, D, G, H) show beech leaf litter mass in the mixed tree species stands (mixed with *Abies* in the North: C, D, and mixed with *Quercus* in the South: G, H, see leaf symbols in the lower right corner of the panels).

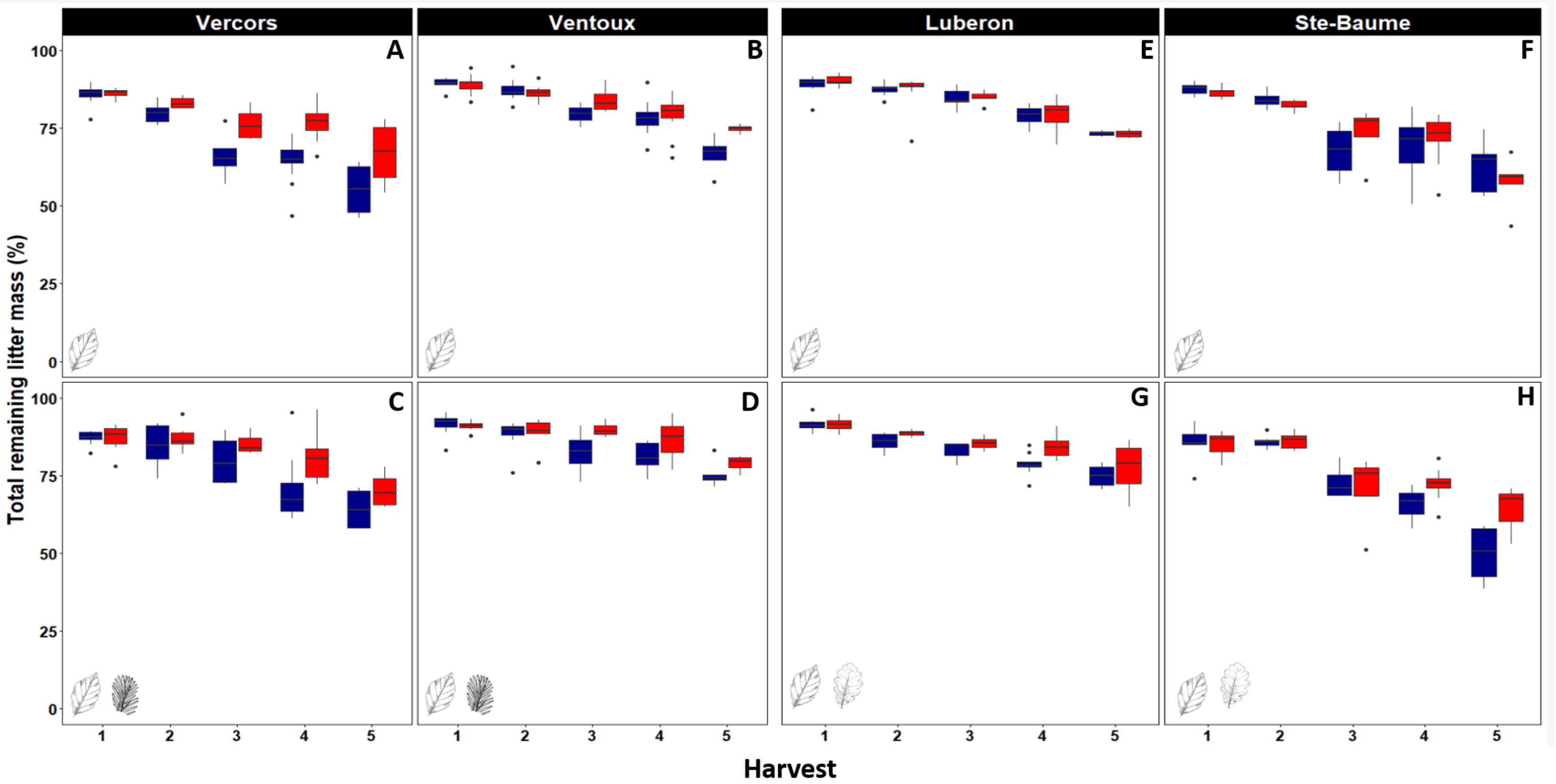

Vercors
Ventoux
Luberon
Ste-Baume
A
B
E
F
C
D
G
H
100
75
50
25
0
Total remaining litter mass (%)
1
2
3
4
5
Harvest

Table S2. Effects of *site*, tree canopy composition (*composition*), rain exclusion (*treatment*), and time of harvest on total remaining mass of beech leaf litter. Coefficient estimate, with standard deviation (in brackets), and part of variance explained are written above. Significant effects (p < 0.05 are shown in bold).

| **Remaining mass ~** | **Estimate** | **% variance explained** |
|---|---|---|
| **Intercept** | **0.86 (+/-0.01)** | **81** |
| **Site** | | |
| *Luberon* | **0.1(+/-0.01)** | **2** |
| *Ventoux* | **0.08(+/-0.01)** | **2** |
| *Vercors* | 0.02(+/-0.01) | - |
| **Canopy composition** | | |
| *Fir-beech* | **0.04(+/-0.01)** | **2** |
| *Quercus-beech* | 0.008(+/-0.01) | - |
| **Treatment** | **0.03(+/-0.01)** | **0** |
| **Time** | **-0.0003(+/-0.00001)** | **7** |
| **Composition: treatment** | | |
| *Fir-beech* | 0.005(+/-0.02) | - |
| *Quercus-beech* | 0.002(+/-0.01) | - |
